# AIE-nanoparticle assisted ultra-deep microscopy in the *in vivo* mouse brain under 1300-nm excitation

**DOI:** 10.1101/2020.12.25.424420

**Authors:** Dong-yu Li, He-qun Zhang, Lina L. Streich, Ping Lu, Ling Wang, Robert Prevedel, Jun Qian

**Affiliations:** Britton Chance Center for Biomedical Photonics, Wuhan National Laboratory for Optoelectronics, Huazhong University of Science and Technology, Wuhan, Hubei, China; State Key Laboratory of Modern Optical Instrumentations, Centre for Optical and Electromagnetic Research, College of Optical Science and Engineering, International Research Center for Advanced Photonics, Zhejiang University, Hangzhou, China; Cell Biology and Biophysics Unit, European Molecular Biology Laboratory (EMBL), Heidelberg 69126, Germany; MoE Key Laboratory for Biomedical Photonics, Huazhong University of Science and Technology, Wuhan, Hubei, China; Zhejiang University Interdisciplinary Institute of Neuroscience and Technology, The Second Affiliated Hospital, School of Medicine, Zhejiang University, Hangzhou, Zhejiang; Candidate for Joint PhD degree from EMBL and Heidelberg University, Faculty of Biosciences, Heidelberg 69126, Germany; State Key Lab of Supramolecular Structure and Materials, Jilin University, Changchun 130012, China

## Abstract

Aggregation-induced emission nanoparticles serve as promising fluorescence probes for multi-photon excitation microscopy due to the large absorption cross-section at NIR-IIb region. Here we present organic AIE nanoparticles that feature high aborption cross-section under three-photon excitation. We show that these enable ultra-deep NIR-IIa excited three-photon imaging in the in-vivo mouse brain.

## Introduction

Modern optical imaging techniques have attracted wide attention in the field of biomedical research due to their key advantages of high resolution and low invasiveness, and thus provide a powerful tool for *in vivo* monitoring of biological structure and function.^1–3^ For example, fluorescence imaging can be utilized to visualize neural activity, vasodilation and -constriction, and blood oxygenation in brain.^4–6^ However, while many optical techniques, including basic fluorescence imaging, permit imaging at superficial layers of tissues, imaging deeper into mammalian tissues is challenging.^4, 7, 8^ While confocal microscopy obtains high 3D resolution, the attenuation of tissue to visible (fluorescence) light caused by absorption and scattering limits the depths of traditional confocal imaging to only a few dozens of microns. Compared to confocal microscopy, two-photon fluorescence microscopy (2PM) significantly extends the imaging depth by employing near infrared-I (NIR-I, 700-900 nm) excitation light. Here, the non-linear excitation confines the fluorescence excitation and thus permits detection of scattered as well as ballistic signal photons. Multi-photon excitation thus enables imaging from a few hundred microns to a millimeter below the surface of scattering, mammalian tissues.^9–13^ To further increase the achievable imaging depth, three-photon fluorescence microscopy (3PM) has been explored and developed in the past.^14–16^ 3PM uses NIR-II (900-1700 nm) excitation light,^17, 18^ which leads to reduced scattering and overall attenuation and thus to increased penetration depth in tissue. Furthermore, the fluorescence of three-photon excitation falls off as~1/z^4^ (where z is the distance from the focal plane), which yields less out-of-focus background and therefore better signal-to-background ratio (SBR) at significant tissue depths.^19^ Despite these clear advantages, 3PM places stringent requirements on the fluorescence probes in terms of high three-photon absorption cross section, quantum yield, photostability as well as biocompatibility. These render most traditional fluorescence probes such as proteins or small molecule dyes challenging for practical 3PM. Therefore, there exists an urgent need to develop novel probes that meet those requirements and are optimized for 3PM and its NIR-II excitation.^20, 21^

Due to their easy modification and remarkable biocompatibility, numerous organic molecules with high quantum yields have been used as fluorescent probes in biological imaging in recent years.^22^ However, most of the organic dyes suffer from aggregation-caused quenching (ACQ) effect due to their planar shaped molecular structure, leading to reduced fluorescence when in the (nano)aggregated state or at a relatively high molecular concentration.^23, 24^ The ACQ effect not only limits the brightness of such molecules for biological imaging, but also restrict their photostability, because organic fluorophores in isolated state at molecular level are more easily photobleached under laser excitation,^25,26^ especially at the ultra-high pulse-energy femtosecond (fs) laser excitation which is typically used in 3PM. ^14–19^

In contrary to the ACQ effect, aggregation-induced emission (AIE) phenomenon can lead to high amounts of fluorescence in the aggregate or solid state.^27–29^ This phenomenon exists in propeller shaped organic molecules which are non-emissive or provide weak fluorescence in benign solvents, but provide orders-of-magnitude higher fluorescence in aggregated form. To exploit this effect, thousands of AIE molecules can be doped into one nanoparticle (NP) to improve their overall absorption cross section, quantum yield and photobleaching resistance,^30^ suggesting AIE-NPs to be excellent three-photon fluorescent probes. In addition, due to the structural flexibility, AIE fluorogens (AIEgens) can in principle be designed for large multiphoton absorption cross sections. For instance, the three-photon absorption cross section of TBDTT is 1.92×10^−81^ cm^6^ s^2^ at 1600 nm^31^, DCDPP-2TPA is 2.95×10^−79^ cm^6^ s^2^ at 1550 nm,^32^ and TPATCN can reach 5.77×10^−79^ cm^6^ s^2^ at 1550 nm,^33^ which is comparable to those of bright quantum dots (QDs)^14, 34^, and far higher than those of common fluorophores, such as fluorescein and GFP (~10^−83^ cm^6^ s^2^).^35^ Accordingly, AIEgens have shown promising performance in high-resolution deep-tissue imaging when excited by three-photon.^31–33^

It is generally thought that the region of NIR-IIb (1500-1700 nm) is more suitable than NIR-IIa (1300-1400 nm) for deep-tissue imaging, because of lower scattering.^36–38^ However, overall tissue absorption is rather strong in the NIR-IIb window because of the water’s absorption spectrum (Fig. 1),^15, 39^ which can cause severe heating-related photodamage and thus limit the maximum excitation laser power for physiological imaging. Altogether, we reason that NIR-IIa excitation might be optimal for *in-vivo* deep-tissue imaging applications^40–42^ despite the slightly enhanced scattering in this wavelength window.

**Fig. 1.**
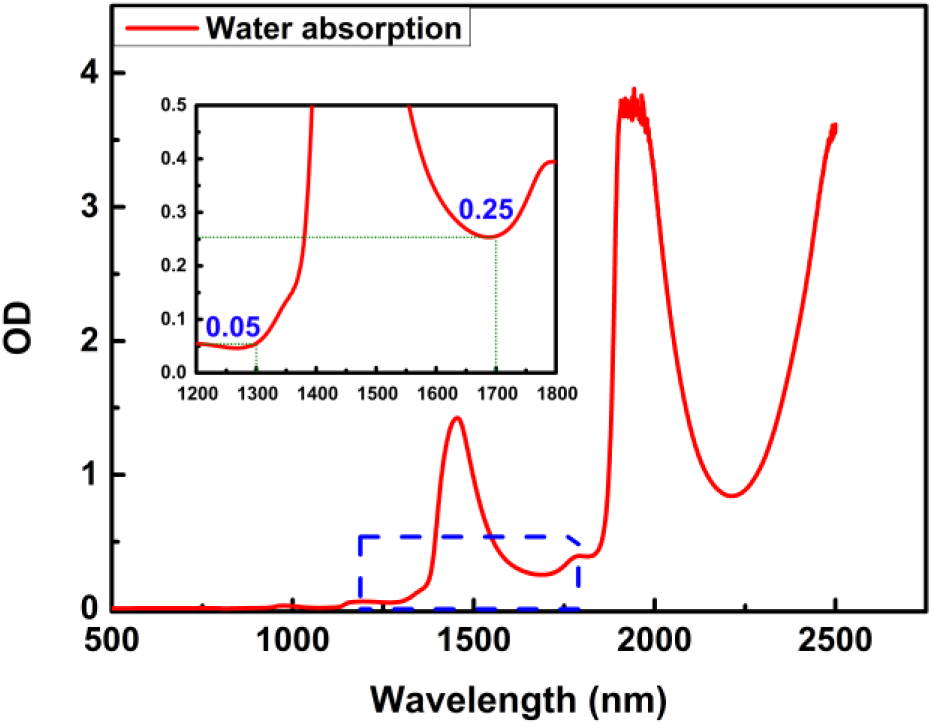
Absorption spectrum and attenuation of water from 500 nm to 2500 nm. The water depth was set to 2 mm, corresponding to the working distance of typical physiology objectives. Inset shows expanded view of blue dashed box area.

In this work, we investigated the performance of AIE NPs optimized for NIR-IIa excitation as three-photon fluorescent probes in deep-tissue imaging. In particular, we chose TPATCN molecules which have previously been shown to exhibit a high three-photon absorption cross section at 1550 nm^33^. After doping them into organic NPs we characterized them for NIR-IIa 3P-excitation at 1300nm. Our results show that the three-photon absorption cross section of TPATCN at 1300 nm is actually increased compared to 1550 nm. Based on this we demonstrated the capabilities of our TPATCN-based AIE NPs for deep-tissue brain imaging. Utilizing a custom-built three-photon microscope, we obtained high signal-to-background ratio (SBR) and an overall imaging depth of ~1.4 mm when imaging sub-cortical vasculature in the in-vivo mouse.

## Results and discussion

### Preparation and optical characterization of TPATCN NPs

As illustrated in Fig. 2, the TPATCN@F-127 NPs were synthesized via a modified nanoprecipitation method which resulted in a hydrophobic TPATCN molecules core and a amphiphilic Pluronic F-127 molecule matrix. The NPs could be dispersed clearly and stably in water, and the overall method was optimized for simplicity and reliability.

**Fig. 2.**
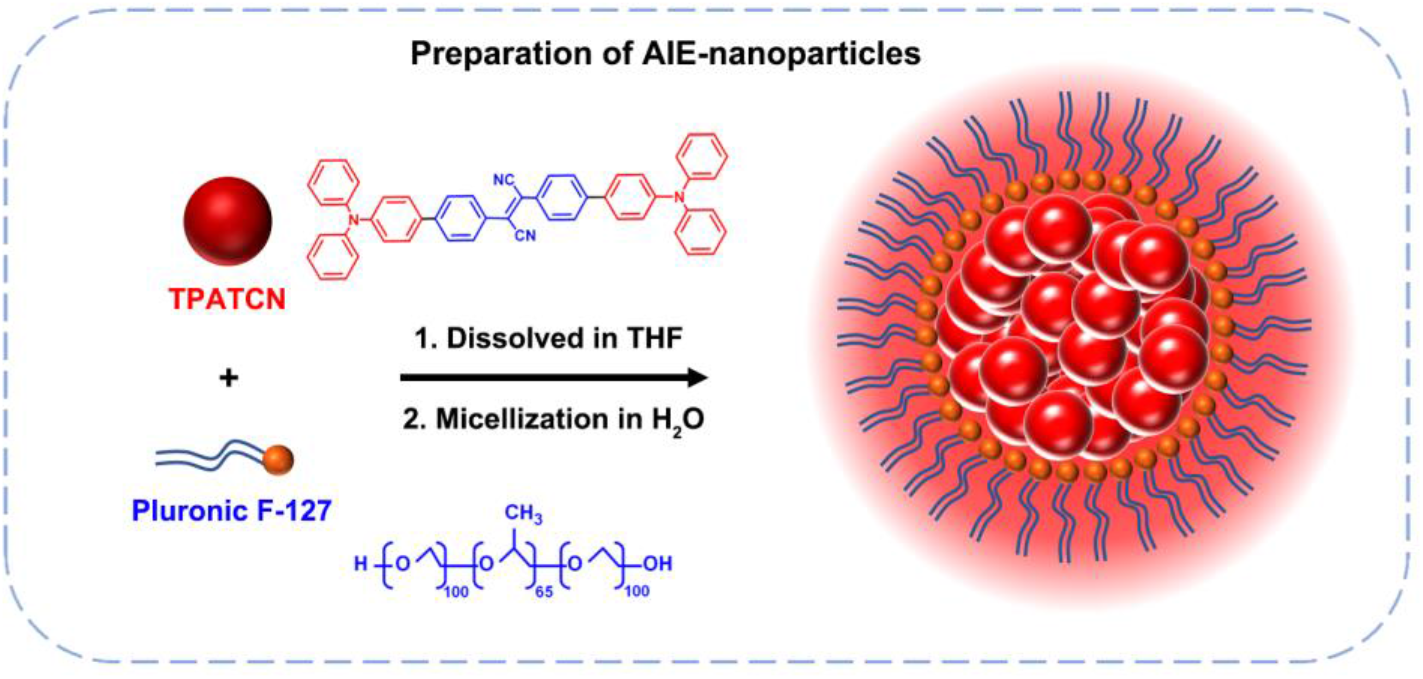
Chemical structures of TPATCN and Pluronic F-127, and schematic illustration of the synthesis of TPATCN@F-127 nanoparticles.

Fig. 3a shows the absorption (blue line) and one-photon fluorescence spectra (red line) of TPATCN@F-127 NPs in aqueous dispersion. We found the absorption peak of TPATCN@F-127 NPs at 444 nm with deep-red fluorescence generated at a peak wavelength of 648 nm.

**Fig. 3.**
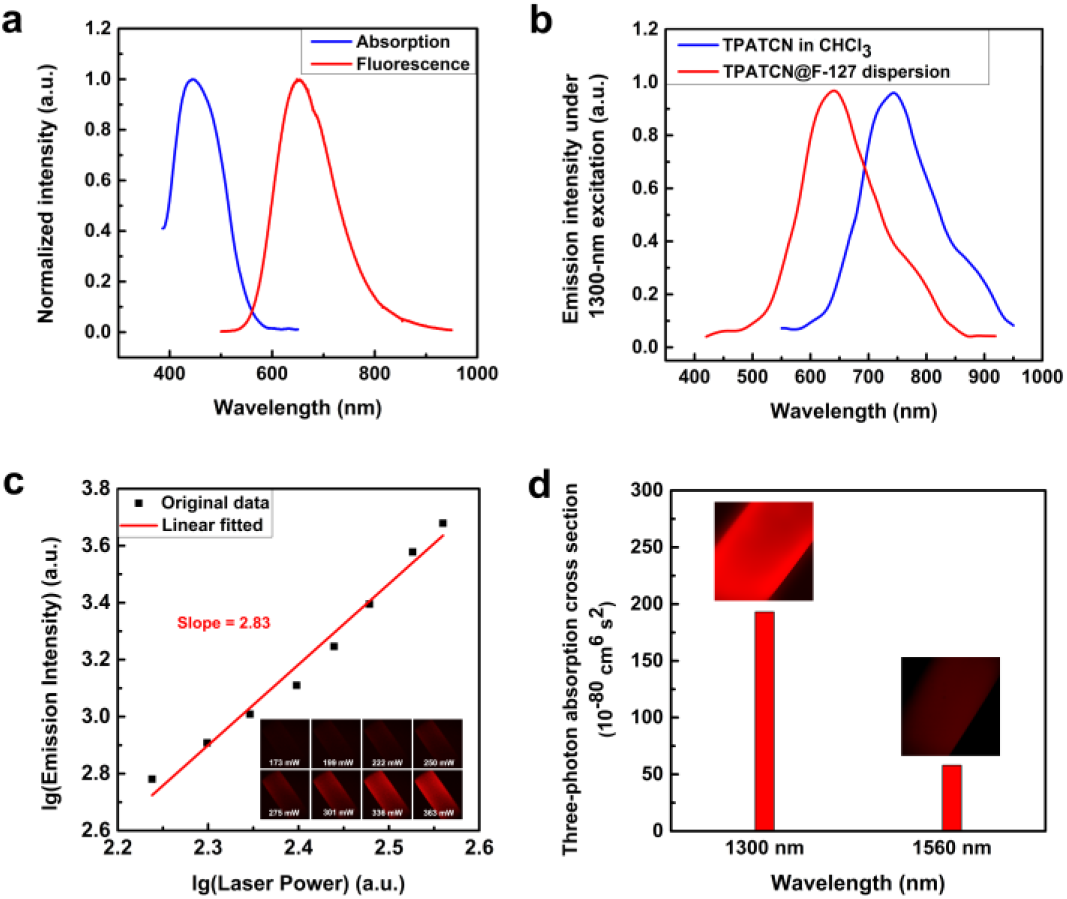
(a) absorption and one-photon fluorescence spectra of TPATCN@F-127 NPs in aqueous dispersion. (b) Emission spectra of TPATCN in CHCl_3_ solution and TPATCN NPs in aqueous dispersion under 1300-nm fs excitation. (c) Relationship of excitation power and fluorescence from TPATCN@F-127 NPs under 1300-nm excitation. The insets are the 3PM images of a glass capillary filled with TPATCN NPs in aqueous dispersion with various excitation power. The power was measured before passing through the microscope optics. (d) Three-photon absorption cross sections at 1300 nm and 1550 nm. The insets are the 3PM images a glass capillary filled with TPATCN NPs in aqueous dispersion under the excitation of 1300 nm and 1560 nm with the same pulse length (200 fs), repetition rate (400 kHz) and average power (20 mW after objective), respectively.

The nonlinear optical characters of TPATCN@F-127 NPs were measured under 1300-nm excitation. As shown in Fig. 3b, both TPATCN in CHCl3 solution (purple line) and in NPs dispersed in water (red line) could generate fluorescence under the excitation of 1300-nm (fs) laser. The emission peak of the NPs was observed at 640 nm, which is effectively unaltered compared to one-photon fluorescence. Compared to the NPs, TPATCN in CHCl_3_ solution generated NIR-I fluorescence under the same fs-laser excitation, with a peak wavelength of 744 nm. We reason that this red shift is due to the smaller polarity of CHCl_3_.

The excitation power dependence relationship of the fluorescence from TPATCN@F-127 NPs was measured to identify the nonlinear emission under the excitation of 1300 nm. The NPs in aqueous dispersion was filled in a capillary glass tube and further imaged by a home-built microscopy system. As shown in Fig. 3c, the fluorescence intensity was rapidly enhanced with the increase of laser power. The logarithm of the fluorescence intensity of TPATCN NPs showed a good linear relation to the logarithm of the excitation power, with a slope of 2.83, which is close to 3. Taking the emission spectrum into account, this suggests that three-photon fluorescence should be the main nonlinear optical process under 1300-nm fs excitation.

Next, we studied the three-photon absorption cross section of TPATCN. Its three-photon absorption cross section has been measured in previous work to be 5.77×10^−79^ cm^6^ s^2^ at 1550 nm.^2^ This allowed us to calculate the three-photon absorption cross section at 1300 nm via comparison of the fluorescence intensity under excitation of these two wavelengths with the same power, pulse length and repetition rate. As shown in Fig. 3d, the three-photon fluorescence was much stronger at 1300-nm excitation than that of 1550-nm excitation, and the three-photon absorption cross section at 1300 nm was thus inferred as 1.93×10^−78^ cm^6^ s^2^, which is the highest value among reported organic dyes, making it a promising optical probe for NIR-IIa excited 3PM.

In addition to the good non-linear optical characters, the TPATCN@F-127 NPs have been reported to have high chemical stability (the absorption and emission spectra remained unchanged in various pH values), high photostability (the fluorescence remained stable under the 1550-nm fs laser irradiation) and high biocompatibility (the injection of the NPs won’t cause inflammation or abnormalities on their major organs, which had experienced the accumulation and excretion of the NPs).^28^ Taken together, TPATCN@F-127 NP are an ideal probe for *in vivo* deep-tissue 3PM under NIR-IIa excitation, which has potentially low photo-toxicity to the animal.

### NIR-I excited *in vivo* deep-tissue 3PM using TPATCN NPs

In order to perform 3PM, we utilized a self-built dual channel laser scanning microscope equipped with a tunable fs laser as shown in Fig. 4. The light source consisted of a 1040-nm fs pump laser and a noncollinear optical parametric amplifier which was tuned to 1300nm excitation wavelength for this study. To maintain short, 50 fs pulses at the sample plane, we built a single-prism pulse compressor.^43^ This ensured a high peak pulse energy which is critical for high 3P excitation efficiency. In this work, only the red channel was used.

**Fig. 4.**
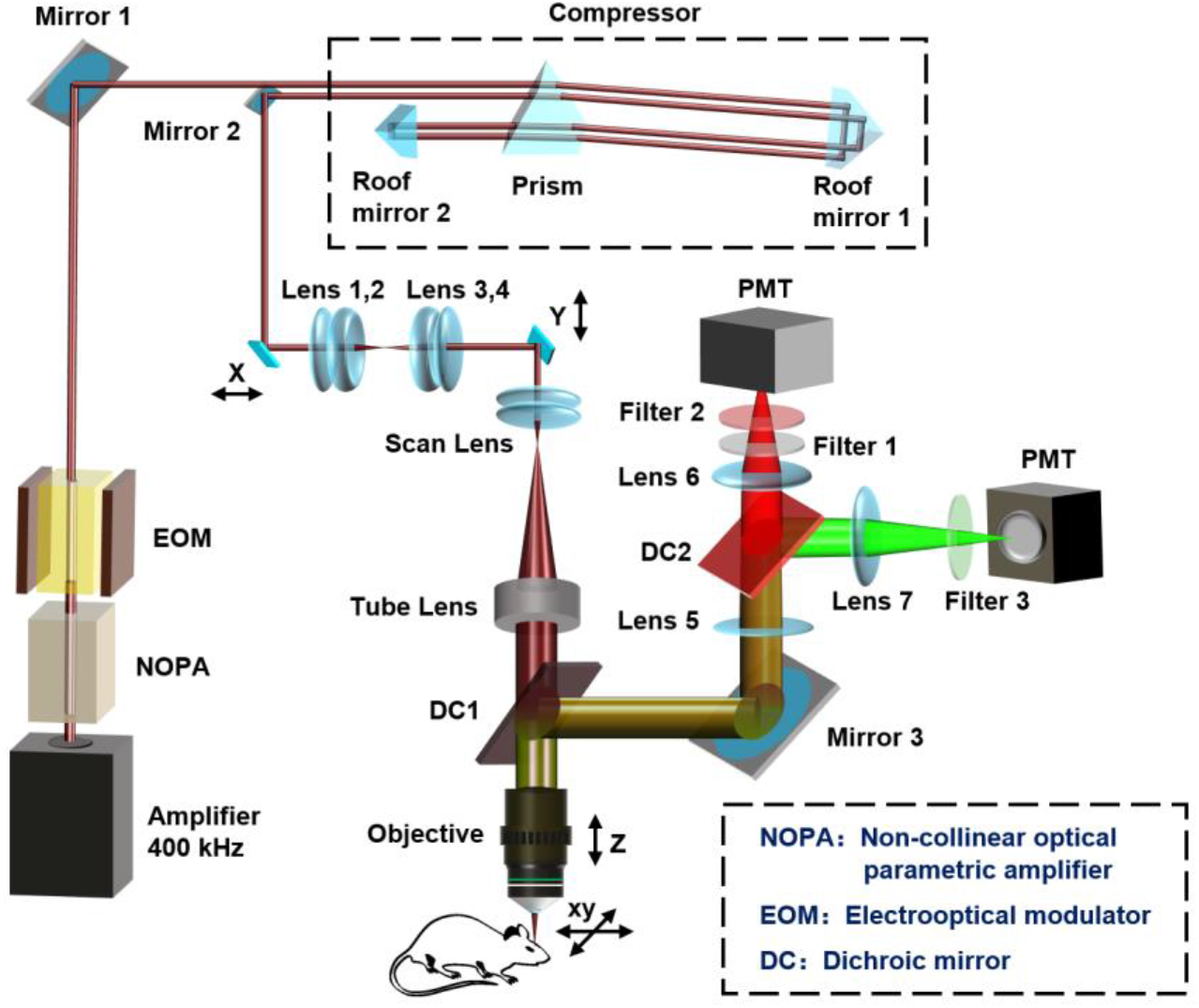
Schematic illustration of the three-photon microscopy system.

Fig. 5 shows the results of TPATCN@F-127 assisted *in vivo* brain 3PM. Due to the promising absorption cross section and biocompatibility, the signals of the NPs were bright and uniform in the vasculature. The unique properties of the NPs enabled high-contrast and deep tissue imaging of the brain vasculature from the surface into deep cortical and sub-cortical regions. The 3PM allowed us to resolve individual microvasculature with diameter as small as a few micrometers even in sub-cortical brain tissue, i.e. below the highly scattering corpus collosum layer. The high SBR enabled 3D vessel reconstruction to depth up to 1360 μm, which represents the deepest in NIR-IIa excited *in vivo* 3PM at this excitation wavelength to date.

**Fig. 5.**
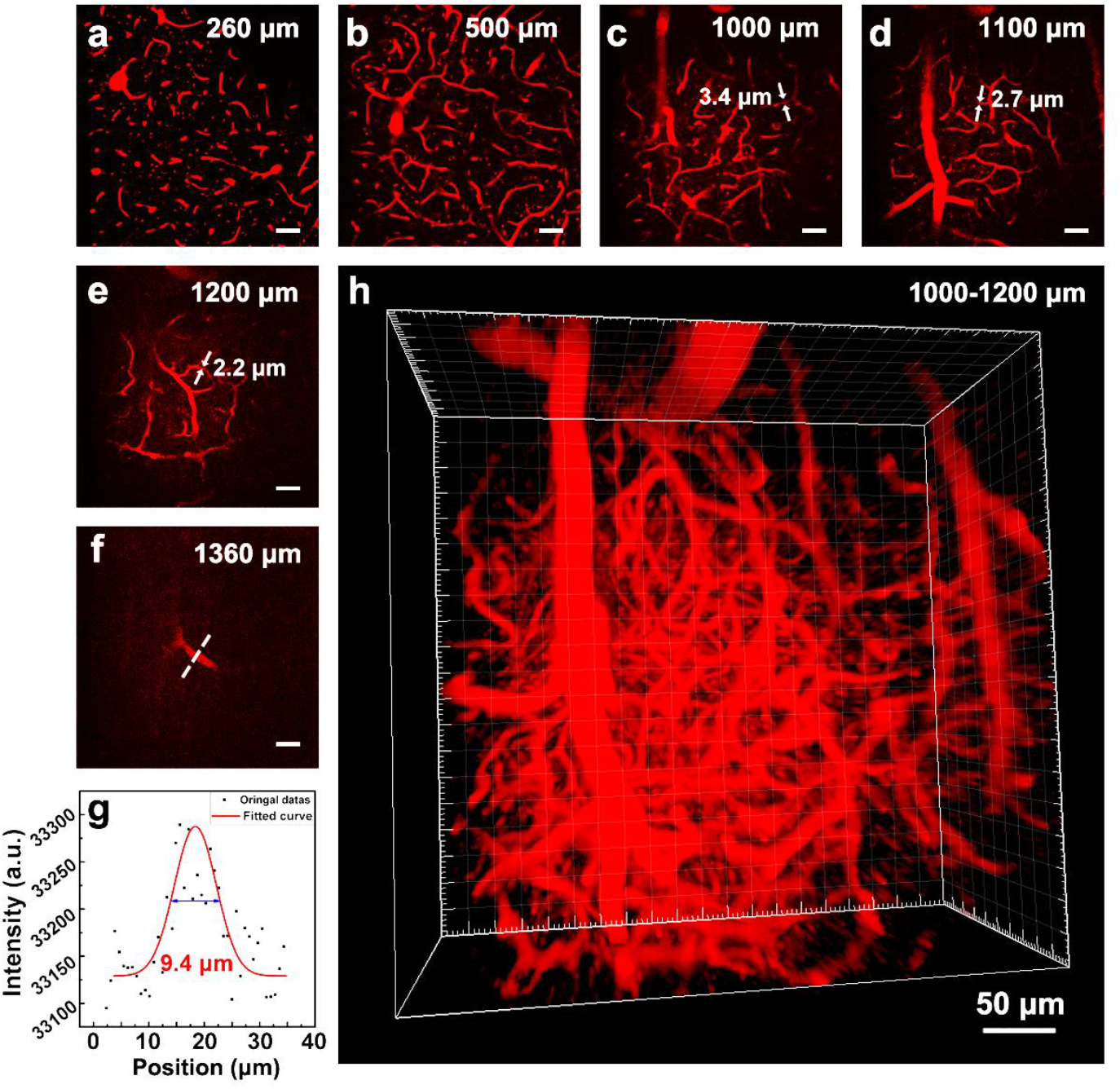
(a)-(f) 3PM images at various brain depth. The values beside the white arrows in (c)-(e) represent the measured diameters of blood vessels (g) Line plot of dashed line in (f). (h) 3D reconstruction of cortical vasculature in the depth range from 1000-1200 μm.

The SBRs at different depths are shown in Fig. 6. The exceptionally high three-photon absorption cross section and QY of TPATCN@F-127 NPs yielded an excellent SBR of up to 24 with only 3.5 mW of laser power after the objective. With the same excitation power, the SBR remained high (>10) until ~500 μm after which the excitation laser power was continuously increased (to a maximum of 35 mW) to maintain the signal intensity. Doing so maintained SBRs of ~10 up to 1000 μm, thanks to the low background associated with 3P excitation. Below 1000 μm, the SBR started to drop due to the limited remaining excitation power at this tissue depths. Nevertheless, the SBR at 1360 μm was still measured to be 2.6. Even deeper imaging (down to SBR ~1) would thus have been possible theoretically, but practical limitations associated with the craniotomy prevented us from lowering the objective further.

**Fig. 6.**
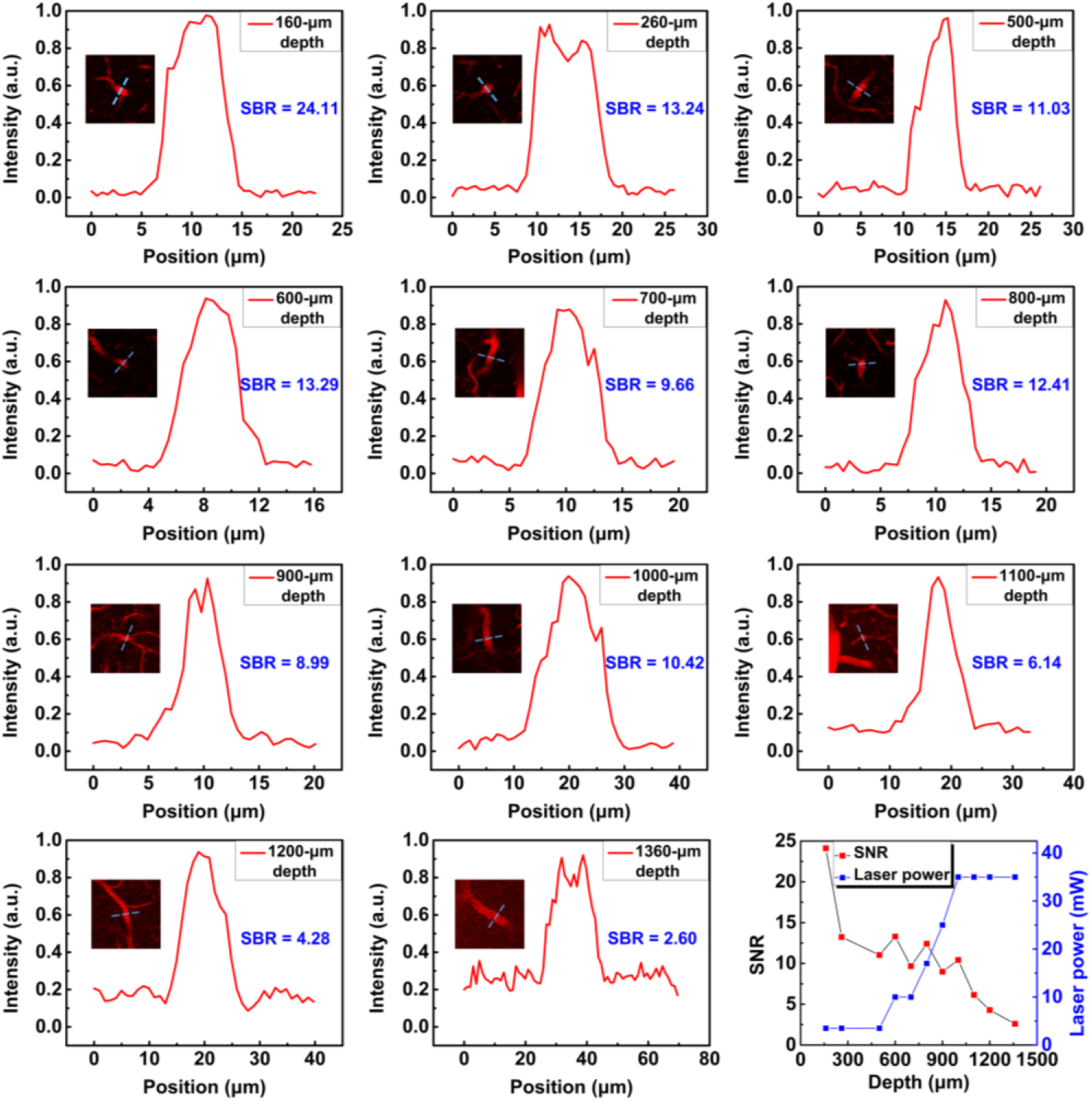
Representative SBR of *in vivo* 3PM at each imaging depth of brain. Insets show the ROIs used for SBR calculation (see dashed lines).

Deep-tissue 3PM relies on high pulse energies to excite three-photon fluorescence. Specifically, the peak pulse power after the objective was as high as 1.75 MW (average power = 35 mW, pulse width = 50 fs, pulse frequency = 400 kHz) when imaging below 1000 μm. Here the comparatively low tissue, i.e. water, absorption of 1300 nm compared to other NIR wavelengths (e.g. 1700nm - see Fig. 1) was crucial in order to prevent photodamage and heating effects which could alter the physiology of the sample. In our experiments, we continuously monitored the breathing and heartbeat of the mouse and found no abnormalities during the imaging process.

## Conclusions

AIEgens are emerging and promising fluorescence probes for 3PM for *in vivo* deep-tissue imaging, due to their high three-photon absorption cross section, high QY in NPs, high photobleaching resistance and high biocompatibility. However, previous work employed excitation lasers exclusively in the NIR-IIb region, where tissue scattering is minimal but water absorption cannot be ignored. Since 3PM requires high, MW-scale peak pulse powers, such NIR-IIb excitation might introduce nonnegligible heating-related photodamage in live animal. Compared to NIR-IIb, NIR-IIa is much more favourable in this respect and can thus be argued to be more biocompatible. As AIE probes optimized for NIR-IIa excited 3PM have not been reported, in this work, we utilized a typical AIE molecule, TPATCN doped into NPs, and explored its capabilities for three-photon microscopy probes. We found that the TPATCN@F-127 NPs could emit bright three-photon fluorescence under the excitation of 1300-nm (NIR-IIa) fs laser. Remarkably, the three-photon absorption cross section at 1300 nm was measured to be as high as 1.93×10^−78^ cm^6^ s^2^, which is the highest among the organic materials reported in the literature. We demonstrated the capabilities of our TPATCN NPs by performing *in vivo* deep-tissue 3PM experiment. Here the NPs exhibited excellent performance and the brain’s microvasculature could be clearly resolved up to a depth of ~1.4 mm, amongst the best reported in the literature. In particular, the SBR remained remarkably high in sub-cortical regions of more than 1 mm depth. Because of the low tissue absorption at 1300nm, no changes in blood vessel morphology, respiration or heartbeat were observed during the imaging process despite the high pulse power of the excitation laser. Thus, TPATCN NPs assisted NIR-IIa excited 3PM holds great potential for sub-cortical brain investigation with low heating-related photodamage.

## Experimental section

### Materials and instruments

The TPATCN were synthesized via the reported protocol. Other chemicals were purchased from Sigma-Aldrich Co., Ltd. The absorption spectra were recorded from 400 to 2500 nm with a spectrophotometer (Cary 5000, Agilent Technologies Inc., USA), at room temperature. The one-photon fluorescence spectrum was recorded using a fluorescence spectrophotometer (F-2500, HITACHI, Japan) with a xenon lamp (excitation range, 300-800 nm).

### Synthesis of TPATCN@F-127 NPs

TPATCN@F127 NPs were synthesized through a modified nanoprecipitation method.^25, 44^ Briefly, 1 mg TPATCN molecule and 10 mg F-127 were solved in 1 mL of THF. The THF solution was then added into 1 mL DI water drop by drop with stirring. Next, the solution was evaporated using a rotary evaporator for 2 min at room temperature. Since THF has a lower boiling point than water, this step could remove the THF, while the water was remained. In this way, TPATCN@F127 NPs in water dispersion was obtained. Similarly, TPATCN@F127 NPs in D_2_O could be obtain by replacing DI water with D_2_O during the process.

### Nonlinear optical spectra measurement

A 1300 nm-fs laser (200 fs, 400 kHz) was focused on a cuvette, which contained TPATCN in CHCl_3_ solution or TPATCN@F-127 NPs in D_2_O dispersion, via a lens (focal length: 5 cm). The laser was generated from an optical parametric amplifier (Orpheus) pumped by a 1040 nm PHAROS-10W fs laser (10 W), in which the emission wavelength can be tuned in the range of 1040-2600 nm. The nonlinear optical emission was collected from the laser-incident direction by an objective (20 × 1.05 NA) and recorded with an optical fiber spectrometer (PG 2000, Ideaoptics Instruments) after passing through a short-pass filter.

### Measurement of power dependence and three-photon absorption cross section

The power dependence and three-photon absorption cross section were measured with an upright scanning microscope (BX61 + FV1200, Olympus) equipped with the laser used in the experiment for nonlinear optical spectra measurement.

To measure the excitation power dependence relationship of the fluorescence from TPATCN@F-127 NPs under 1300-nm excitation, a capillary containing TPATCN@F-127 NPs in D_2_O dispersion was imaged via the microscopy system. The excitation wavelength was set at 1300 nm, and the images were captured with various excitation power (the output powers were set as 173 mW, 199 mW, 222 mW, 250 mW, 275 mW, 301 mW, 336 mW and 363 mW). Then, the relationship between fluorescence intensity and excitation power was calculated.

To measure the three-photon absorption cross section, a capillary containing TPATCN in CHCl_3_ solution was imaged via the microscopic system. The images at 1300-nm excitation and 1550-nm excitation with the same power were captured. The three-photon absorption cross-section of TPATCN was calculated according to the following equation:^45^

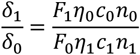

where δ is the three-photon absorption cross section, F is the three-photon fluorescence intensity, η is the QY, c is the molar concentration, n is the refractive index of the solvent, and the subscripts 0 and 1 represent the reference (at 1550 nm) and the sample (at 1300 nm). Herein, η_0_=η_1_, c_0_=c_1_, n_0_=n_1_. Therefore, the formula could be simplified as follow:

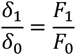

where δ_0_ is 5.77×10^−79^ cm^6^ s^2^.

### Three-photon fluorescence microscopic system

For the 3PM of the mouse brain, we used a home-built laser scanning microscope and a 1300 nm fs laser (maximal output power 400 mW, 400 kHz, 35 fs) from a noncollinear optical parametric amplifier (Spectra Physics) pumped by a regenerative amplifier (Spirit-16, Spectra Physics). We used a home-built dispersion compensation unit to maintain short pulse durations (50 fs) at the sample plane. The maximum laser power after the objective was measured to be 35 mW. Detection included proper filters (FF01-593/LP-30-D & F01-940/SP-30-D, Semrock) and PMT (H107740, Hamamatsu). Data acquisition was based on ScanImage.^46^

### Animals

All animal experiments were approved by EMBL’s Institutional Animal Care and Use Committee. 8-week old female C57 mice were used for in vivo 3PM imaging of brain vasculature.

### In vivo three-photon microscopy of cortical vasculature of mice

To obtain optical access to the brain, a cranial window was implanted as previously described.^47^ During surgery animals were anesthetized with 5% isoflurane vapor mixed with O_2_ for induction and maintained at 1-1.5%. For cranial window preparations, a 7mm diameter craniotomy was made over the motor cortex, centered at 0.6 mm posterior and 0.7 mm lateral to the Bregma point, and a 7-mm coverslip (~170 μm thick) was placed on top of the brain.

Before imaging experiments, The TPATCN@F-127 NPs in PBS (200 μL, 1 mg/mL) were injected into the circulatory system of the mice through the tail vein. Then, the mice were fixed and put under the microscope.

The image stack was acquired with a depth interval of 5 μm, and pixel dwell time was 10 μs, 5 frames were averaged. The exciation power was exponentially increased with depth and the maximum average power under the objective was 35 mW.

During imaging experiments, mice were anesthetized with isoflurane (2% in oxygen, Harvard Apparatus) and positioned onto a small animal physiological monitoring System (ST2 75-1500, Harvard Apparatus), which allows to maintain animal body temperature at 37.5°C. During experiments eyes were covered with eye ointment.

## Data quantification

All imaging data were analyzed with Image J software that was developed by National Institutes of Health (Bethesda, Maryland). 3D image reconstruction was performed in Imaris (Bitplane).

## Acknowledgement

This work was supported by National Natural Science Foundation of China (NSFC) (Grant Nos. 61735016, 82001877 and 61975172); China Postdoctoral Science Foundation funded project (Nos. BX20190131, 2019M662633); Zhejiang Provincial Natural Science Foundation of China (LR17F050001); Funding for Postdoctoral Innovation Research Post in Hubei Province. Fundamental Research Funds for the Central Universities (2020-KYY-511108-0007). Work at the EMBL was supported by the European Commission (EU grant FET-PROACTIVE, No. 951991 ‘Brainiaqs’). The authors also thank Dr. Senthilkumar Deivasigamani for the surgeries and Prof. Cornelius Gross for support.

